# The Invisible Maze Task (IMT): Interactive Exploration of Sparse Virtual Environments to Investigate Action-Driven Formation of Spatial Representations

**DOI:** 10.1101/278283

**Authors:** Lukas Gehrke, John R. Iversen, Scott Makeig, Klaus Gramann

## Abstract

The neuroscientific study of human navigation has been constrained by the prerequisite of traditional brain imaging studies that require participants to remain stationary. Such imaging approaches neglect a central component that characterizes navigation -the multisensory experience of self-movement. Navigation by active movement through space combines multisensory perception with internally generated self-motion cues. We investigated the spatial micro genesis during free ambulatory exploration of interactive sparse virtual environments using motion capture synchronized to high resolution electroencephalographic (EEG) data as well psychometric and self-report measures. In such environments, map-like allocentric representations must be constructed out of transient, egocentric first-person perspective 3-D spatial information. Considering individual differences of spatial learning ability, we studied if changes in exploration behavior coincide with spatial learning of an environment. To this end, we analyzed the quality of sketch maps (a description of spatial learning) that were produced after repeated learning trials for differently complex maze environments. We observed significant changes in active exploration behavior from the first to the last exploration of a maze: a decrease in time spent in the maze predicted an increase in subsequent sketch map quality. Furthermore, individual differences in spatial abilities as well as differences in the level of experienced immersion revealed an impact on the quality of spatial learning. Our results demonstrate the feasibility to observe behavioral changes associated with spatial learning, opening the way to the study of cortical dynamics of navigation.

## 1 Introduction

Access to some form of mental spatial representation is a prerequisite for successful navigation in known environments. Here, we introduce a new experimental paradigm, the Invisible Maze Task (IMT) for the study of navigation in which freely-moving participants must compute and use a spatial representation by relying on interactively triggered sparse visual or other sensory feedback defining the virtual walls of a maze. The maze is wholly invisible, and stimuli defining the walls are only transiently revealed as the participant reaches out to ‘touch’ the virtual walls. Place yourself in the following scenario: waking up in a dark and unknown hotel room in the middle of the night, you need to visit the toilet. By reaching and touching the walls you manage to make your way without locating the light switch. On the way back, you contemplate whether to turn on the light or whether you had gained sufficient spatial knowledge to safely return to bed by confirming your internal spatial representation with a few select wall touches. This example demonstrates our ability to use egocentric (body-centered, self-to-object relations) sensory information to compute a so-called allocentric spatial representation of the environment sufficiently detailed and accurate to guide future action. Egocentric tactile feedback from wall touches combined with vestibular and proprioceptive information as well as motor efference copies is used to compute a spatial representation of the room and corridor layout to allow navigation back to bed. The IMT captures the richness of such navigation, revealing in a tractable manner the multi-modal aspects involved in real-world navigation. Real-world navigation has been difficult to study precisely because it involves a wide variety of information about location and orientation in space derived from self-motion cues that include motor efference copies, movement-related sensory information originating inside the body, and flow signals experienced in the auditory, visual and somatosensory modalities to successfully build complex spatial representations. The requirement of free movement has generally precluded the addition of brain measurement modalities. For example, in prior EEG neuroimaging experimental protocols to investigate the neural basis of spatial cognition, movement-related information is absent largely for fear of movement-induced artifacts. Limitations of other established imaging methods also do not allow participant movement because of physical constraints of the sensor [16, 17, 19, 29]. As a consequence, neuroscientific studies investigating natural human behavior during free-roaming dynamic interactions with external environments are relatively rare. Knowledge about how and where the human brain processes sensory cues to form spatial representations, and about factors influencing individual’s performance in spatial cognitive tasks, has been derived predominantly from stationary experiments [23, 28]. Beyond greater realism, another argument in favor of an ambulatory paradigm is that animal studies have provided substantial new insights into the neural representations of location and direction in physically moving animals [30, 45]. In providing evidence of selectively firing neuronal populations across several brain regions encoding spatial information about location, i.e. place-, grid-and boundary vector cells, and heading direction, these studies significantly contributed to the understanding of the neural representations of spatial cognition in settings with higher ecological validity, see [7, 9] for a comprehensive overview. Meanwhile, studies with human participants, i.e. classical passive viewing and cursor-controlled paradigms, are missing key components of the *natural* subjective experience of space.

### 1.1 Formation of Spatial Representations through Action-Oriented Percepts

A well-established theory of spatial learning in children assumes an ontogenetic sequence from egocentric to allocentric (external world-centered, object-to-object relations) representations of space implying a sequential development from coarse/simple to complex spatial representations [20, 31, 36]. In this framework, the first stage of spatial knowledge entails encoding sensory representations of landmarks. In the second stage, route knowledge develops through repetitive travel (and/or mental rehearsal of travel) along one or more routes between previously encountered landmarks. The final stage involves connection of different routes within a map-like model of the environment. This defines so-called survey knowledge, an allocentric spatial representation allowing planning of new routes, shortcuts, and detours. It is reasonable to assume an underlying continuum in the micro genesis of spatial knowledge as compared to a strictly categorical spatial learning [3, 15]. For instance, presumably allocentric systems evaluating sensory spatial signals are active in parallel, e.g. place, grid and boundary cells in the rat brain [30, 45] and process inputs continuously. Taken together, survey knowledge, ultimately conceived in an allocentric reference frame, develops from egocentric representations of idiothetic spatial signals driving the buildup of metric representations, i.e. turn angles and distances [4, 5], over repeated travels. Idiothetic signals generated at turns highlight and specify the association between the place in the environment and the action taken [4]. Prior findings indicate that bottom-up spatial micro genesis through active exploration of unknown environments is inherently ego-dependent [10].

### 1.2 A new approach: The Invisible Maze Task

To allow investigation of behavior as well as human brain dynamics reflecting spatial learning in freely ambulating humans, we developed a virtual reality paradigm in which participants learn the spatial layout of mazes by active exploration. In this Invisible Maze Task (IMT), we task subjects to explore mazes by touching otherwise invisible walls to receive sensory feedback (visual, auditory or other) about the location of the wall. At the end of the exploration phase, participants have to produce a bird’s eye view map representation that presumably requires mental transformations of egocentrically experienced spatial signals into allocentric representations. After exploring and building a survey representation of a previously unknown environment, the resultant spatial model can be tested through egocentric sampling of spatial information when the same environment is experienced again. This learning and confirming strategy of spatial representations highlights the interplay of egocentrically perceived information and an allocentric representation [10]. Hence, participants never experience an externalized, i.e. completely visible, representation of the entire maze structure but have to rely on internally generated spatial knowledge to complete the task. In the following, we describe the paradigm based on *visually* sparse, interactive virtual exploration and describe behavioral parameters that can be extracted to investigate spatial learning. Specifically, we investigated whether self-report as well as psychometric measures in the spatial domain had predictive power explaining the production of allocentric spatial representations over time [22, 38, 43]. Understanding cognition as optimizing the outcome of behavior, we hypothesized changes in body dynamics occurring during formation and consolidation of spatial representations. Therefore, the number of spatial exploratory movements and overall time spent exploring were tested as predictors of the buildup of spatial representations. Specifically, we hypothesized that exploratory movements and time spent in mazes would be reduced as spatial representations become more accurate, possibly as a consequence of optimization of the energy costs of querying the spatial environment.

## 2 Methods

To test our hypotheses, we captured body motion while participants freely explored an interactive sparse “Invisible Maze” environment by walking and probing for virtual visual wall feedbacks with their hand, delivered by a virtual reality (VR) headset. Participants explored four different mazes in three consecutive maze trials each. At the end of each maze trial, participants were asked to draw a sketch map of the maze from a bird’s eye view perspective as an index of spatial learning.

### 2.1 Subjects

Thirty-two healthy participants (aged 21-47 years, 14 men) took part in the experiment. All participants gave written informed consent to participation and data collection in accordance with the declaration of Helsinki approved by the local ethics committee (proposal number: GR_08_20170428). Three participants were excluded from data analysis due to incomplete data caused by technical issues in two cases and difficulties in complying with the task requirements in one case.

### 2.2 Equipment

Data collection was performed at the Berlin Mobile Brain/Body Imaging Labs (BeMoBIL), a 10 m x 15 m research facility equipped with wireless high-density EEG synchronized to motion capture and virtual reality. Participants interactively explored virtual mazes walking around the lab space wearing a head-mounted display (HMD). Visual stimulation was presented via an Oculus VR (Facebook Inc., Menlo Park, California, USA) Rift DK2 HMD (100° nominal field of view horizontally and vertically, 960 x 1080 pixels per eye, 75 Hz frame rate). Head position and orientation were updated by fusing data from the headset’s internal inertial sensors and using a six-LED rigid body mounted to the headset and tracked via PhaseSpace (PhaseSpace Inc., San Leandro, California, USA) Impulse X2 system. To update the head position, motion capture data was sampled at 240 Hz and smoothed by averaging across one frame update of the HMD, approximately 13.3 milliseconds. To correct orientation drifts originating from unstable inertial data, we continuously calculated an offset between the stable orientation of the motion capture rigid body and the unstable magnetometer data. A difference exceeding 3° in the Euler yaw direction triggered a correction in all three Euler dimensions by 1° per second until the difference approached 0°. Four further rigid bodies consisting of four LEDs each were attached to the lower arm, upper arm and both feet. To update the position of the right hand in VR, positional data from a PhaseSpace glove with 8 LEDs was smoothed by averaging across one frame update. Visual stimuli were generated on a MSI (MSI Co. Ltd, Zhonghe, Taiwan) Gaming Laptop (MSI GT72-6QD81FD, Intel i7-6700, Nvidia GTX 970M) using Worldviz (Santa Barbara, California, USA) Vizard Software worn in a backpack. Participants were further equipped with a microphone and headphones for audio communication and masking of auditory orientation cues. For EEG data collection, a 160 channel wireless BrainProducts MOVE System (Brain Products GmbH, Gilching, Germany) was administered with 128 channels applied on the head and 32 channels on the neck.

### 2.3 Psychometric and Self-Report Measures

After arrival to the lab, participants were given a number of questionnaires and self-report measures to complete, in order to characterize individual experience with virtual reality, individual differences in spatial abilities and preferred navigational styles:

**Perspective Taking and Spatial Orientation Test, PTSOT** Participants viewed an array of objects on a sheet of paper and by taking the perspective of one of the objects judged the angle between two other objects in the array and sketch it in [27]. We recorded the absolute deviation from the correct angle.

**Santa Barbara Sense of Direction Scale (Freiburg Version), FSBSOD** This questionnaire is a measure of self-ascribed navigational ability consisting of 15 items [21]. We took the average of all correctly recoded items as the final measure.

**Igroup’s Immersion and Presence Questionnaire, IPQ** The IPQ measures the sense of presence experienced in a virtual environment (VE). We processed the results according to [35].

**Gaming Experience** We asked participants to indicate how long they have been playing video games and further asked them to rate their gaming skills. A composite measure taking the sum of the two standardized scores was calculated as the final measure.

**Reference Frame Proclivity Test, RFP** This online available tool determines the proclivity of participants to preferentially use either an egocentric or an allocentric reference frame during a virtual path integration task [14,18]. For further correlation analysis, allocentric reference frame proclivity was coded as “1”, egocentric as “0” and a tendency to switch reference frames as “2”.

**Simulator Sickness Questionnaire, SSQ** The SSQ measures simulator sickness on three factors: nausea, oculomotor and disorientation [25]. We administered SSQ twice, before and after the experiment and used the average value of each sub-scale as the resulting measure.

### 2.4 Sparse Virtual Environments

Four different environments consisting of invisible virtual walls, 90° turns and a starting point were defined. All paths were composed of ten 1x1m squares spatially arranged to different layouts (I, L, Z, U; see figure 1 B). Participants were instructed to explore the space by walking and reaching in order to probe the walls to either side of the paths (see figure 1 A). Upon collision of the hand with an invisible wall, a white disc was displayed 30cm behind the collision point parallel to the invisible wall (see figure 1 C). The disc grew in size as participants moved their hand further into the wall and reached its maximum size (30cm diameter) when the participant’s hand was 30cm deep into the wall. After two seconds the disk faded out. Another second later the disc reappeared colored red. The visual feedback was reset after participants retreated their hand back out of the wall. Participants were instructed to repeatedly touch the walls and not leave their hand *in the wall* to avoid the disc turning red.

**Fig. 1.**
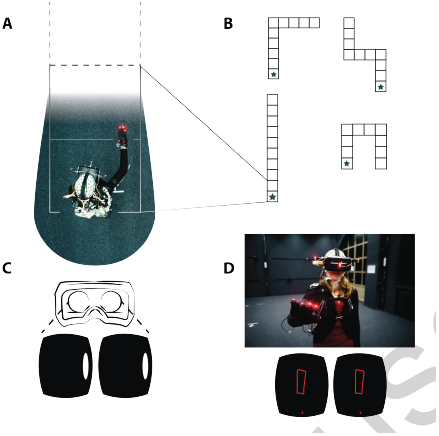
**A** Participant displayed from a bird’s eye view located at the starting point of an “I”-maze. The star marks the starting position but was not visible during the experiment. Participants were instructed to explore the maze and return to the start after full exploration of the maze. **B** Four mazes were used in the study including an “I”, “L”, “Z”, and “U” shaped maze clockwise from lower left to lower right. Each maze was explored three times before the next maze was learned. **C** Exemplary first-person view in *binocular* “VR optics” of subject in A (above) touching the wall to the right. **D** Top: after returning to the starting location, participants drew a top-down view of the explored maze. The participant wears high-density wireless EEG, head-mounted virtual reality goggles and LEDs for motion capture attached to the hands, goggles, and torso. Bottom: screenshot of drawn sketch map. As a visual guidance during drawing, a small red dot was rendered at the position of the tracked hand. This figure is licensed CC-BY and available on Figshare [13].

### 2.5 Procedure

The complete experiment, including EEG preparation took approximately 4 hours. Participants first completed the questionnaires and self-report measures. Subsequently, the EEG electrodes were prepped. Next, participants were instructed to explore a path until the end, i.e. reaching a dead end, and subsequently find their way back to the starting position. Participants were informed that touching a wall with the right hand to either side of their body would temporarily illuminate a small part of the wall. Further, the participants were briefed that they will explore the same path three times in a row and that there will be four different paths to explore over the entire experiment. For all twelve trials, participants were oriented in the direction of the corridor by the experimenter after a short disorientation phase where participants were led walking in circles. A gamified feedback was displayed reporting back the performance in the current maze trial. The feedback measure was based on the progress made in the current path and provided information whether the end of a path was reached and how many wall collisions of the hands and the head were registered.

#### Drawing Task

After each exploration phase, participants were asked to draw a map of the environment. The experimenter entered the lab space at the end of each trial and handed the participant a computer mouse and asked them to use it to trace their mental map of the environment just explored. Participants were instructed to start drawing by clicking once with the left mouse button. A red sphere appeared in the VR goggles at the tracked position of the right hand (see figure 1 D). Holding down the left mouse button, participants were able to draw a red line by moving their hand in space. Finally, participants were instructed to take a camera screenshot of their drawing by pressing down the mouse wheel once and holding their final drawing in view. Participants were allowed to erase their drawing and restart at any time by pressing the right mouse button.

### 2.6 Data Processing & Statistical Analysis

#### Motion capture measures

For quantification of changes in exploration behaviors we extracted the following measures for the time window between the start of each maze exploration and the return to the initial square: (1) the total number of wall touches, (2) the total duration of the exploration, and (3) the average velocity of participants in the maze (the first derivative of raw positional data of the head rigid body in the (x,z) ground plane).

#### Sketch map Measure of Spatial Ability

Two independent raters judged each sketch map drawing of each maze trial per maze and subject. A total number of 29 x 12 sketch maps were rated with respect to their usefulness as a navigational aid. The raters were presented with the question:“Imagine that you can take the present sketch map with you into the virtual environment and use it as a navigational aid. How useful would the map be for you?” To give their rating between 0 (= no help at all) and 6 (= very helpful) they were given the correct shape of the maze to be rated side by side with the drawing to rate [2]. To test inter-rater reliability, we computed Cohen’s Kappa with squared weights to emphasize larger rating differences using R (R Development Core Team), Version 3.4.3, and package *irr* [6, 12].

#### Trial Rejection

Prior to the final statistical analysis, the data was cleaned. First, trials were rejected when more than ten wall collisions with the head were recorded. This resulted in the rejection of six out of a total of 348 trials. Second, four individual trials were rejected due to procedural problems during data collection. In one case, the battery of the LED driver of the motion capture system died, whereas in the remaining case a LED cable became loose and the trial had to be aborted. Lastly, we deemed all trials incomplete where the path was not fully explored. With this criterion, 26 trials were rejected. Overall, 312 trials remained, amounting to 89.7 % of the total number of trials.

#### Linear Mixed Effects Model

To investigate changes in exploration behavior, we performed linear mixed effects analyses of the relationship between (1) maze trials and (2) maze configurations and each dependent measure “number of wall touches”, “duration” as well as “movement velocity” using R package *lme4* [32]. As fixed effects we entered “maze trial” with three repetitions for each maze and “maze” with four levels (different mazes) as well as their interaction. As random effects we considered intercepts for participants as well as by-maze trial random slopes for the effect of each dependent variable “number of wall touches”, “duration” as well as “movement velocity”. Subsequently, to examine changes in the sketch map drawings, we fit identical linear mixed effects models to the dependent measure sketch map usefulness. Ultimately, to make inferences about the relationship between the three measures of body dynamics on sketch map usefulness, we fit linear mixed effects models to the dependent variable “sketch map usefulness”. As fixed effects we entered the body movement measures “number of wall touches”, “duration” as well as “movement velocity”. As random effects we considered intercepts for participants as well as by-maze trial random slopes for the effect of each sketch map usefulness. We assumed an underlying continuum in the sketch map ratings “usefulness” and hence conceived the variable as interval scaled. For all analyses, we obtained P-values by calculating likelihood-ratio tests of the full model with the effect in question against the model without it [42]. Post-hoc, we tested non-parametric pairwise differences using uncorrected Wilcoxon signed-rank tests [40]. We considered significant results with *α <* 0.05. For report generation and data visualization purposes we used R packages *knitR, ggplot2, ggpubr, cowplot* and *corrplot* [24, 37, 39, 41, 44].

#### Correlation of Psychometric and Self-Report Measures

We preprocessed all questionnaire and self-rating data to construct a full correlation matrix including dummy coding of binary variables. We added gender as an additional binary factor of interest with female participants coded “0”. For better understanding, scores of perspective taking (PTSOT) were recoded, so that large numbers indicated better performance. Finally, we created a correlation matrix with the average ratings of the two sketch map raters together with the motion capture measures and the subjective data.

## 3 Results

First, we investigated if our experimental manipulation changed the exploration behavior of participants. Therefore, we tested if body movement measures were explained by repeated maze trials or changes in the maze configurations.

**Duration** The repeated measures factor maze trial affected the duration between start and end of each maze exploration (*χ*^2^(2)=15.521, p<0.001) lowering it by about 31.7 seconds ± 8.1 (standard errors) from the first to the second maze trial and 39.9 seconds ± 9.4 (standard errors) from the first to the third maze trial. Subsequent non-parametric pairwise comparisons revealed clear reductions in exploration times for the comparison of the first and second maze trial (p=0.09), the first and third maze trial (p=0.05), with a diminished reduction between the second and third maze trial (p=0.47) (see figure 2 A). Different maze configurations also affected the duration of maze exploration (*χ*^2^(3)=11.109, p<0.05). Initial exposure as well as increasingly complex maze configurations were associated with increasing exploration times (see figure 3 A). The interaction of both factors revealed an effect of maze trials on exploration duration (*χ*^2^(6)=12.333, p=0.05).

**Fig. 2.**
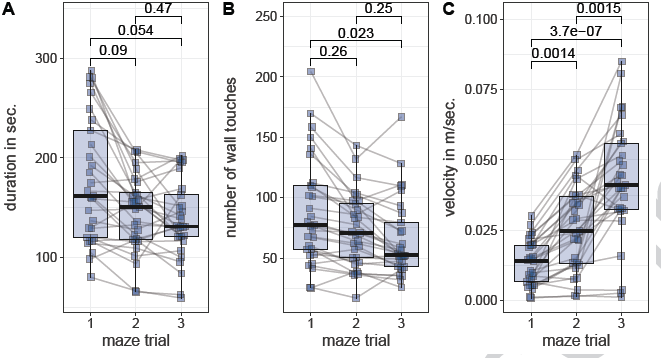
Box-Whisker plots with individual observations of each participant averaged across maze configurations for each repeated maze trial 1 to 3. **A** Duration in seconds elapsed between the start and end of each exploration phase, **B** Number of wall touches during the exploration phase and **C** Movement Velocity in meters per second. P-values of pairwise comparisons are calculated by non-parametric Wilcoxon signed-rank tests.

**Fig. 3.**
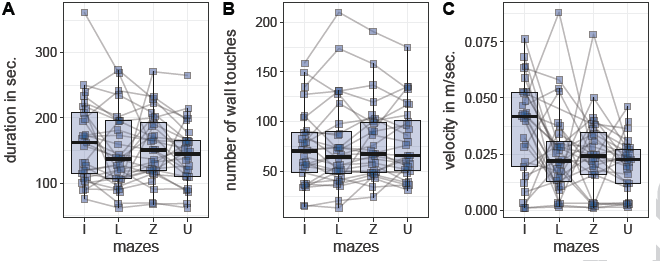
Box-Whisker plots with individual observations of each participant averaged across maze trials exploring each maze configuration “I”, “L”, “U” and “Z”. **A** Duration in seconds elapsed between the start and end of each exploration phase, **B** Number of wall touches during the exploration phase and **C** Movement Velocity in meters per second.

**Number of Wall Touches** The number of wall touches was significantly af-fected by repeated measurements across maze trials (*χ*^2^(2)=21.37, p<0.001) with a reduction in the number of wall touches by 15 touches 3.8 (stan-dard errors) from the first to the second maze trial and 22 touches 4.1 (standard errors) from the first to the third maze trial. Neither the factor maze (*χ*^2^(3)=3.94, p=0.27) nor the interaction of both factors (*χ*^2^(6)=4.32, p=0.63) revealed an impact on the number of wall touches. Non-parametric pairwise comparisons between maze trial were not significant for the comparison of first and second maze trial or between second and third maze trial However, between the first and third maze trial (p<0.05) the number of touches decreased significantly (see figure 2 B).

**Movement Velocity** Participants’ movement speed was significantly increased across maze trials (*χ*^2^(2)=29.495, p<0.001) with an increase by 0.01 m/sec ± 0.003 (standard errors) from the first to the second maze trial and 0.03 m/sec ± 0.004 (standard errors) from the first to the third maze trial. Participants’ movement speed was also affected by maze (*χ*^2^(3)=22.513, p<0.001) with a significant decrease after exploration of the “I” maze. We registered no interaction effect. Post-hoc multiple comparisons were significant for all comparisons between the three maze trials (1-2: p<0.01, 2-3: p<0.01, 1-3: p<0.001) and between mazes “I” and “L” (p<0.05), “I” and “Z” (p<0.05) and “I” and “U” (p<0.01), and did not show the same attenuation effect in later maze trials seen for maze duration. As an index of the consistency of this finding, all but one participant (28 of 29) moved faster in the third maze trial as compared to the first (see figure 2 C).

### 3.1 Sketch Map Ratings and Changes in Body Movement Behavior

Cohen’s *Κ* was calculated with squared weights of rating differences to determine if there was agreement between two raters judgments on the usefulness of a given sketch map to navigate a virtual environment. A total of 312 maps were rated. There was very high agreement between the two raters judgements, *Κ* = 0.835, *p <*0.001. Next, to test for a general effect of the experiment exposure on the sketch map ratings a Wilcoxon signed rank test was tested with the mean sketch map rating after the first trial run against the null hypothesis of a mean equal to zero. A deviation from 0 for the ratings of the first sketch maps would indicate that participants successfully built a mental representation of the invisible maze they explored for the first time. We observed a true location of the mean (=3.2) different from 0 (p<0.001) indicating successful spatial learning after the first trial. Investigating changes in the sketch map ratings over the repeated maze trial (*χ*^2^(2)=3.8123, p=0.15) and maze configurations (*χ*^2^(3)=3.2171, p=0.36) as well as their interaction (*χ*^2^(6)=4.3286, p=0.63) revealed no significant impact, however we did observe higher average sketch map ratings for 19 of 29 subjects after the third trial run as compared to the first. To test the predictive power of body movement measures on the usefulness of the sketch maps, a null model was compared to three models each with one additional predictor. We subsequently added “duration”, “number of wall touches”, as well as “movement velocity” as predictors. The duration between start and end of each maze exploration affected the sketch map ratings (*χ*^2^(1)=17.160, p<0.001) as did the number of wall touches (*χ*^2^(1)=3.852, p<0.05). We observed no impact of the head velocity on the sketch map ratings (*χ*^2^(1)=0.7221, p=0.4).

### 3.2 Correlation with Psychometric-and Self-Report Measures

To examine the spatial exploration behavior and sketch map usefulness and their relation with subjective measures, a correlation matrix among all measured variables was calculated. For the sketch map usefulness, we observed significant positive correlations with perspective taking skills (*r* = 0.39, *p* < 0.05) as well as experienced realism inside the VE (*r* = 0.41, *p* < 0.05). We observed a highly significant correlation between gender and gaming experience (*r* = 0.65, *p* < 0.001) with male participants scoring higher on gaming experience. Furthermore, male participants reported a higher general sense of presence (*r* = 0.42, *p* < 0.05) after the VE exposure with all presence sub-scales being correlated. Finally, we found a strong correlation between the duration of maze exploration and the number of wall touches (*r* = 0.5, *p* < 0.01) revealing more explorative touches the longer a maze was explored.

**Fig. 4.**
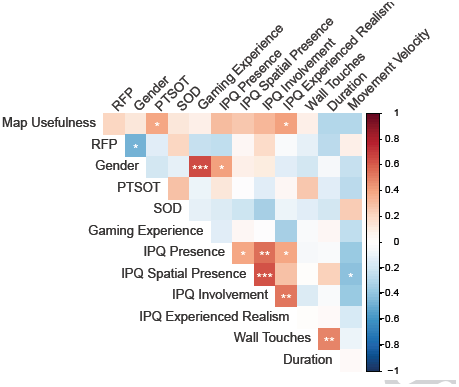
Correlation Matrix among Psychometric-and Self-Report Measures. Each cell is colored as a function of the strength of positive (red) or negative (blue) correlation. Significant correlations are indicated by * (P<0.05), *\*\** (P<0.01), or *** (P<0.001). Concerning sketch map usefulness, we observed significant correlations with PTSOT (*r* = 0.39, *p* < 0.05) as well as high correlations across all four Presence Scales. Furthermore, duration significantly correlated (*r* = 0.5, *p* < 0.01) with the number of wall touches.

## 4 Discussion

We introduced a new paradigm, the Invisible Maze Task, to investigate realistic spatial learning as reflected through the computation and use of spatial representations derived from transient, discrete visual feedback in sparse (virtual) environments. We observed significant changes in body movement behavior across repeated explorations of the same environment. Participants moved faster, for a shorter period of time and with less touches of the surrounding walls supporting the assumption that each transient wall touch event may carry information used for formation and updating of some form of spatial representation. We hypothesize that the sparse environments were learnt in an efficient way that allows optimization of energy costs of repeated future explorations. Therefore, we argue, that the environment was incorporated in a useful way to guide future behavior.

### 4.1 Spatial Learning in Sparse Environments

To quantify the quality of sketch maps we chose the subjective rating measure “map usefulness” because it has been proven useful for within-subject investigation [2]. Although we observed a high variability between participants, 27 out of 29 participants clearly demonstrated spatial learning as reflected in the usefulness of their sketch maps after the initial exposure to the environment. Subsequent changes in the usefulness ratings were only minimal with a slight, albeit non-significant, increase with repeated exposures to the environment. In other words, we found that participants either drew a useful map after the first encounter with an environment and then maintained a level of usefulness in their sketch maps, or they never drew a meaningful map at all. In line with previous findings [1, 22], we noticed a substantial increase in sketch map quality over time only in a very few participants. The fact that some participants were able to draw a near perfect representation of the explored environment without ever witnessing full vista space may be interpreted against a linear, sequential development of spatial knowledge [36] that should have been observable through incremental improvements of the drawn sketch maps usefulness. To investigate a potential connection between spatial exploration behavior and sketch map quality we fitted linear mixed effects models of the spatial exploration parameters number of touches, times in a maze, and movement velocity to the sketch map ratings. The biggest impact on sketch map usefulness originated from the time spent exploring a maze with a decrease in duration predicting an increase in sketch map quality. In addition, a decrease in wall touches predicted higher sketch map quality. We conclude, that brief exploration phases are sufficient for most of the participants to form and maintain a spatial representation through discrete and localized samples of an otherwise invisible environment. Wall touches during repeated exposure to the same environment then serve as a probing mechanism for the correctness of the internal representations, just like finding your way back to bed in the middle of the night by confirming the location of walls and doors. This change in behavior reflects a more efficient means to navigate conserving energy when moving through a known environment. By administering several established measures of navigational abilities we investigated the impact of individual differences in spatial abilities and preferences on spatial learning in the invisible maze task. Contrary to previous findings we observed no impact of gaming experience on scores of the SOD, PTSOT and the spatial sketch mapping task [33]. However, most findings on the interplay between immersion, spatial presence, involvement and spatial abilities stem from experiments with 2-D displays. The recent surge of affordable head-mounted VR technologies will shed light on how accurate these new technologies map a three-dimensional reality. In the current study, we observed covariations between the feeling of presence experienced during exploration and the usefulness of the resulting sketch maps. One possible explanation for this correlation is that participants with a higher immersion score were able to better hold a realistic representation of the environment in memory and were subsequently better able to draw it. Two widely used metrics of spatial abilities are the psychometric perspective taking and orientation test and the sense-of-direction scale [21, 27]. PTSOT scores were significantly correlated with sketch map usefulness ratings. We instructed subjects to draw a top-down map moving their hand in the air imagining they were drawing on a chalkboard. Therefore, perspective taking processes, i.e. changing the viewpoint, were engaged during drawing of an accurate sketch map. A high correlation between perspective taking ability and sense-of-direction provides the ground for future group-based analysis approaches of good and bad spatial learners. Interestingly, we observed no substantial correlation of sense-of-direction with any other measure of interest. Finally, we investigated the impact of individual reference frame proclivities [14, 18] on the performance in the invisible maze task. In an online test, participants were classified into three groups reflecting a preference for egocentric reference frames, a preference for allocentric reference, or the flexible switch between reference frames. We did not find any significant correlation of the preferred reference frame with spatial learning in the IMT. It is reasonable to assume that reference frame proclivities as measured in a passive visual flow paradigm without vestibular feedback do not play any role in a more natural active exploration setting such as the IMT. It is well established that vestibular information is used to update egocentric representations of position and orientation [26, 34]. The absence of any impact of reference frame proclivities as measures with the RFPT again indicates that traditional desktop-based measures of spatial abilities might not reflect behavior in natural three-dimensional environments accurately.

### 4.2 Limitations

In the current study, we used sketch maps as a dependent measure for the quality of internal spatial representations. We used a qualitative assessment instead of quantifiable data, e.g. segment lengths and angular accuracy between segments. This approach is prone to subjective tendencies but was countered in the present study by measuring the agreement between two raters. Still, other sketch map measures might provide a deeper insight into the variables that affect the accuracy of mental spatial representations derived from exploration of the environment. Sketch maps are always subject to individual’s capabilities to draw and their belief in their skill as well as the interpretation of the rater [11]. Furthermore, participants were required to carry a substantial amount of equipment to render our virtual environment. Therefore, participants were restricted, to a certain degree, in their ability to move as they would naturally move without the equipment. We may use more explicit tests of spatial knowledge in future environments, such as asking how many turns it would take to get from one point to another or asking for a bearing to a distant point such as the entrance.

### 4.3 Summary and Future Directions

We introduce the Invisible Maze Task to study spatial learning behavior during ambulatory exploration of sparse environments that provide only transient feed-back, therefore breaking down spatial explorative inputs into discrete moments in time. Our paradigm provides the basis for investigations of “atoms of spatial thought” that ultimately allow computation of spatial representations of an environment that has never been seen as a whole. This approach, which we have behaviorally validated here, will enable investigation of the tight link of physical behavior and cognitive processes during spatial learning. Furthermore, because we simultaneously measure brain activity using EEG, we plan to next use event-related EEG neuroimaging to test existing models of spatial cognition [3] and to gain a deeper understanding of the relationship of cognition, active spatial exploration behavior, and brain dynamics. For example, we plan to investigate oscillatory brain networks underlying computations between egocentric and allocentric representations of space [8]. The results from this initial study offer a new perspective on the interplay between body dynamics and common assessments of spatial orientation skills during the formation of spatial representations.

